# *Proteus mirabilis* employs a contact-dependent killing system against competing *Enterobacteriaceae*

**DOI:** 10.1101/2021.03.19.436238

**Authors:** Dara Kiani, William Santus, Kaitlyn Kiernan, Judith Behnsen

**Affiliations:** Department of Microbiology and Immunology, University of Illinois at Chicago, College of Medicine, Chicago, IL

**Keywords:** *Proteus mirabilis*, inter-bacterial competition, bacterial killing, *Enterobacteriaceae*

## Abstract

Many bacterial species encode systems for interference competition with other microorganisms. Some systems are effective without contact (e.g. through secretion of toxins), while other systems (e.g. Type VI secretion system (T6SS)) require direct contact between cells. Here, we provide the initial characterization of a novel contact-dependent competition system for *Proteus mirabilis*. In neonatal mice, a commensal *P. mirabilis* strain apparently eliminated commensal *Escherichia coli*. We replicated the phenotype *in vitro* and showed that *P. mirabilis* efficiently reduced viability of several *Enterobacteriaceae* species, but not Gram-positive species or yeast cells. Importantly, *P. mirabilis* strains isolated from humans also killed *E. coli*. Reduction of viability occurred from early stationary phase to 24h of culture and was observed in shaking liquid media as well as on solid media. Killing required contact, but was independent of T6SS, the only contact-dependent killing system described for *P. mirabilis*. Expression of the killing system was regulated by osmolarity and components secreted into the supernatant. Stationary phase *P. mirabilis* culture supernatant itself did not kill but was sufficient to induce killing in an exponentially growing co-culture. In contrast, killing was largely prevented in media with low osmolarity. In summary, we provide the initial characterization of a potentially novel interbacterial competition system encoded in *P. mirabilis*.

**IMPORTANCE:** The study of bacterial competition systems has received significant attention in recent years. These systems collectively shape the composition of complex ecosystems like the mammalian gut. They are also being explored as narrow-spectrum alternatives to specifically eliminate problematic pathogenic species. However, many competition systems that effectively work *in vitro* do not show strong phenotypes in the gut. Our study was informed by an observation in infant mice. Further *in vitro* studies confirmed that *P. mirabilis* was able to kill several *Enterobacteriaceae* species. This killing system is novel for *P. mirabilis* and might represent a new function of a known system or even a novel system, as the observed characteristics do not fit with described contact-dependent competition systems. Competition systems are frequently present in multiple *Enterobacteriaceae* species. If present or transferred into a probiotic, it might be used in the future to reduce blooms of pathogenic *Enterobacteriaceae* associated with disease.

## INTRODUCTION

Bacteria frequently inhabit densely populated environments like the soil or the human gastrointestinal tract. In these environments competitive interactions, such as interference and exploitative competition are common (1). In exploitative competition, bacteria compete for common nutrient sources, whereas in interference competition bacteria encode systems that directly affect communication or viability of competitor bacteria. A multitude of different competition systems have been described and are continuing to be discovered (2, 3). They are used for “bacterial warfare” in the gastrointestinal tract by commensal and pathogenic species and are thought to collectively shape the composition of the gut microbiota (4, 5). In the current antibiotic crisis, tools to precision edit the microbiota and to eliminate only problematic species instead of the majority of gut bacteria are highly desirable (1, 6). Narrow-spectrum competition mechanisms encoded by commensals to kill closely related species have therefore received significant attention in recent years (7–11).

Bacterial competition systems are classified as either contact-dependent or contact-independent. Contact-independent competition is mediated through secreted compounds like classical antibiotics, bacteriolytic enzymes, or bacteriocins, colicins, and microcins (12). Contact-dependent mechanisms have been discovered predominantly during the last decade and include, amongst others, Type VI-secretion system (T6SS)-mediated effector translocation (13–16), contact-dependent inhibition (CDI) (17–20), contact-dependent inhibition by glycine zipper proteins (Cdz) (21), and microcin proximity-dependent inhibition (MccPDI) (22). These systems require direct contact between cells, as effector molecules are not diffusing from the producing cell but are transferred when cells touch, e.g. through the molecular syringe complex of the T6SS. Functions of the effector molecules are wide-ranging and include interference with protein synthesis and inducing pore formation in the bacterial membrane of the target cell (12).

*Enterobacteriaceae* are successful colonizers of the mammalian gut, in the form of commensals, pathobionts, or pathogens. They dominate the microbiota in two instances: in the infant gut (23, 24) or during dysbiosis of the adult gut (25). In an undisturbed adult gut environment, *Enterobacteriaceae* levels are generally low (26, 27). One of the most frequent *Enterobacteriaceae* species in the mammalian gut is *Escherichia coli*. However, species from other genera including *Klebsiella*, *Enterobacter*, *Serratia* and *Proteus* can be frequently isolated from both healthy infants and adults (28, 29). Recently, the genus *Proteus* has been reclassified to the new family *Morganellaceae*, forming together with *Enterobateriaceae* the order *Enterobacteriales* (30). For simplicity, we use the old terminology throughout our manuscript. Our study focuses on *E. coli* and *Proteus mirabilis*. *P. mirabilis* is a known pathogen of the urogenital tract but is also frequently present as a commensal in the gastrointestinal tract. However, due to sampling methods, *Proteus* species abundance is often underestimated. *Proteus* species were found in only 7.8% of healthy adult fecal samples (31) but are present in 46% of jejunal and duodenal mucus samples (32). They are more frequently isolated from patients with diarrhea and Crohn’s disease but a direct role in the disease has not yet been established (33). *Proteus* species and *E. coli* were isolated from the majority of cesarean-born infants in Pakistan (34) indicating that *P. mirabilis* and *E. coli* are both found in the human gut. Blooms of commensal *Enterobacteriaceae* are often associated with disease, e.g. inflammatory bowel disease (IBD) or necrotizing enterocolitis in infants (NEC) (35, 36). However, probiotic species of *Enterobacteriaceae* also exist. The probiotic *E. coli* Nissle 1917 is used to treat intestinal diseases like diarrhea in infants (37).

Here, we characterize a contact-dependent competition system employed by *P. mirabilis* to compete with other members of the *Enterobacteriaceae* family. We initially observed the competition between *P. mirabilis* and *E. coli in vivo* in mouse pups and replicated the phenotype *in vitro*. The killing system functions independently of T6SS, the only contact-dependent killing system described for *P. mirabilis*. The system is therefore novel for *P. mirabilis* and might represent a novel system for *Enterobacteriaceae* in general. The focus and scope of this article is the initial characterization of this *P. mirabilis* competition system *in vitro.* Further studies are warranted to reveal the genetic components, regulatory circuits, and *in vivo* importance of this competition system, as well as its prevalence in other species.

## RESULTS

In any natural environment where bacteria reside, such as the gastrointestinal tract (GI), aquatic or soil environments, bacteria will compete with one another in order to survive and replicate. We observed what appeared to be efficient competition between two commensal strains of *Enterobacteriaceae* in mice. A female specific pathogen free (SPF) mouse naturally colonized with *Escherichia coli* was mated with a male SPF mouse naturally colonized with *Proteus mirabilis*. Surprisingly, pups were exclusively colonized with *P. mirabilis* and no *E. coli* was identified in their feces post-weaning (Fig. 1A). We expected *E. coli* and *P. mirabilis* to be transferred to pups through intimate contact and coprophagy. As we recovered only *P. mirabilis* from the pups, we investigated this apparent competitive advantage of *P. mirabilis*.

**Figure 1:**
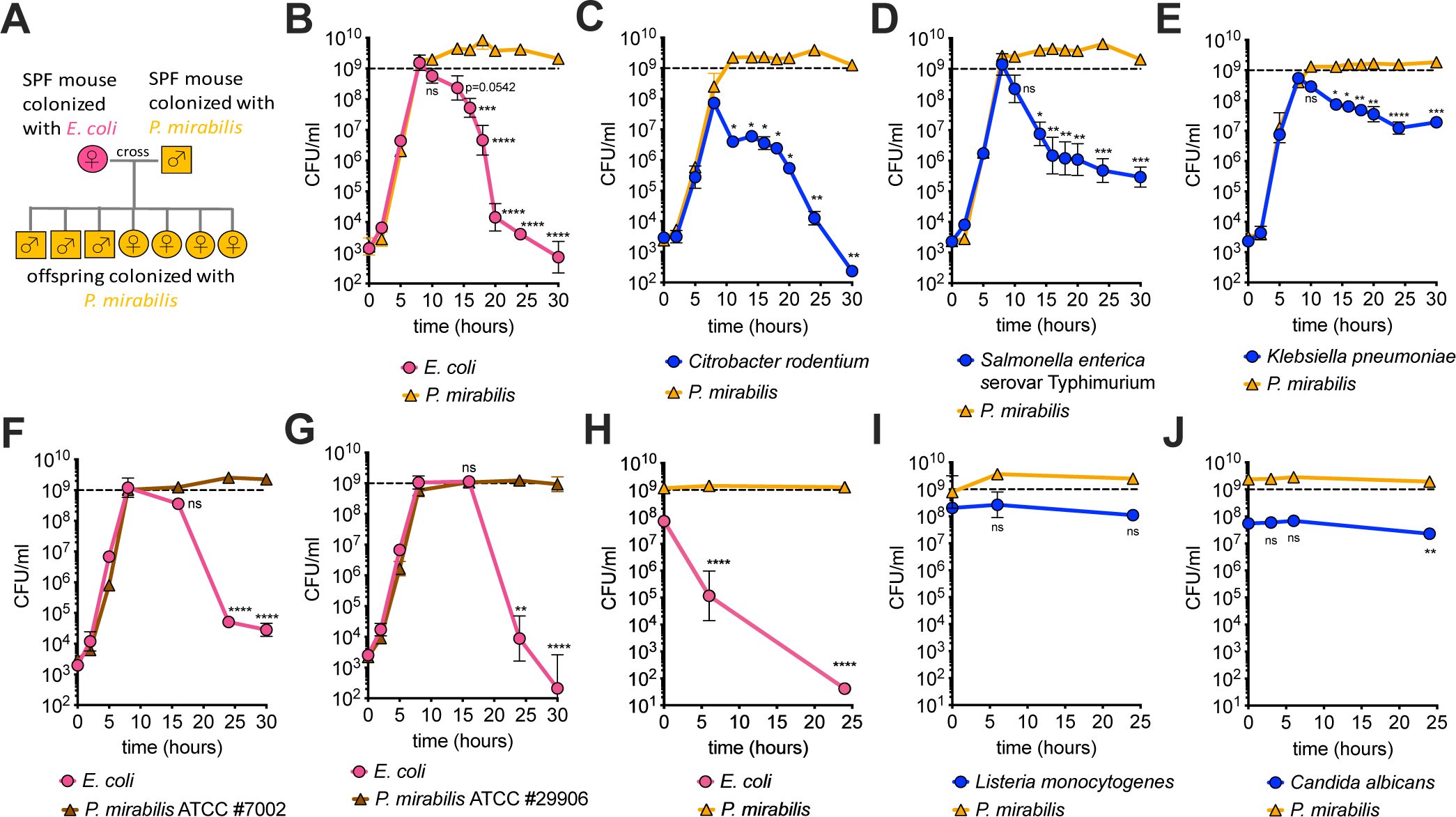
*Proteus mirabilis* reduces the viability of *Enterobacteriaceae*. **(A)** Schematic representation of breeding scheme. The offspring of an SPF female mouse colonized with *Escherichia coli* and an SPF male mouse colonized with *P. mirabilis* were exclusively colonized with *P. mirabilis*. **(B-E)** Shaking liquid growth curve of *P. mirabilis* isolated from mice and different *Enterobacteriaceae* species in co-culture in LB: (**B**) *E. coli*, (**C**) *Citrobacter rodentium*, (**D**) *Salmonella enterica* serovar Typhimurium (**E**) *Klebsiella pneumoniae*. **(F+G)** Shaking liquid growth curve of *P. mirabilis* (**F**) ATCC #7002 or (**G**) ATCC #29906 and *E. coli* in co-culture in LB. **(H-J)** “Killing assay” with *P. mirabilis* and target species (**H**) *E. coli*, (**I**) Gram positive *Listeria monocytogenes,* or (**J**) yeast *Candida albicans. P. mirabilis* to target species ratio was ∼10:1. All data represent mean +/-SEM of at least three biological replicates. If error bars are not visible, error is smaller than symbol size. For panels B through G, a one-way ANOVA was performed comparing the 8-hour time point (upon entry to stationary phase) with the remaining time points (stationary phase). For panels H through J, a one-way ANOVA was performed comparing the 0-hour time point and remaining time points. * = p<0.05, **= p<0.01, ***= p< 0.001, ****= p<0.0001, ns = not significant.

### *P. mirabilis* reduces the viability of *Enterobacteriaceae* species

We serotyped multiple isolates of *E. coli* and *P. mirabilis* from our mouse colony and found only one serotype of *E. coli* (PennState *E. coli* Reference Center, data not shown) and *P. mirabilis* (Vitek2 analysis, data not shown). Different lines of mice were either colonized with *E. coli* or *P. mirabilis*, but not both species at the same time. We grew the isolated *E. coli* and *P. mirabilis* strains in a co-culture in Lysogeny broth (LB). During the exponential phase of growth, both strains grew equally well. In stationary phase, *P. mirabilis* maintained its viability. The number of viable *E. coli* on the other hand declined quickly, reaching a rate of 1600-fold loss of viable cells within 2 hours (Fig. 1B). In a monoculture of *E. coli* we observed no loss of viability in stationary phase (Fig. S1). *P. mirabilis* also significantly reduced the viability of *Citrobacter rodentium,* a rodent pathogen and model organism for EHEC in mice, and the human pathogen *Salmonella enterica* serovar Typhimurium (Fig. 1C, D). In contrast, the viability of another *Enterobacteriaceae* member, *Klebsiella pneumoniae,* was not reduced to the same extent as the others (Fig 1E). The phenotype was independent of capsule, as no increase in susceptibility was seen with a capsule deficient *K. pneumoniae* strain (data not shown). Importantly, the ability to kill extends to other *P. mirabilis* strains. Two isolates of *P. mirabilis* (the type strain and a human isolate) also killed *E. coli* in shaking liquid media (Fig. 1F, G). We next tested if *P. mirabilis* can reduce viability of the Gram-positive bacterium *Listeria monocytogenes* and the eukaryotic yeast *Candida albicans.* Since both species grow poorly in LB, we directly assessed viability by adding prey species to stationary cultures of *P. mirabilis*. *P. mirabilis* drastically reduced the viability of *E. coli* in this assay (Fig. 1H), but we observed no change in the viability of *L. monocytogenes* (Fig 1I) or *C. albicans* (Fig. 1J).

### Killing of *E. coli* is an active process that requires direct contact and live *P. mirabilis* cells

We next wanted to test if *P. mirabilis* killing requires contact or if it occurs via the release of proteins, molecules, membrane vesicles, or phages into the surrounding medium. We found that in cell-free stationary supernatant of *P. mirabilis* (pH ∼7.6), *E. coli* cells remained viable (Fig. 2A). *P. mirabilis* therefore does not seem to reduce viability of *E. coli* through a contact-independent mechanism. Formalin-fixed stationary *P. mirabilis* cells also failed to reduce the viability of *E. coli* in stationary phase supernatant or in fresh LB (Fig. 2B,C). Killing therefore requires live *P. mirabilis* cells.

**Figure 2:**
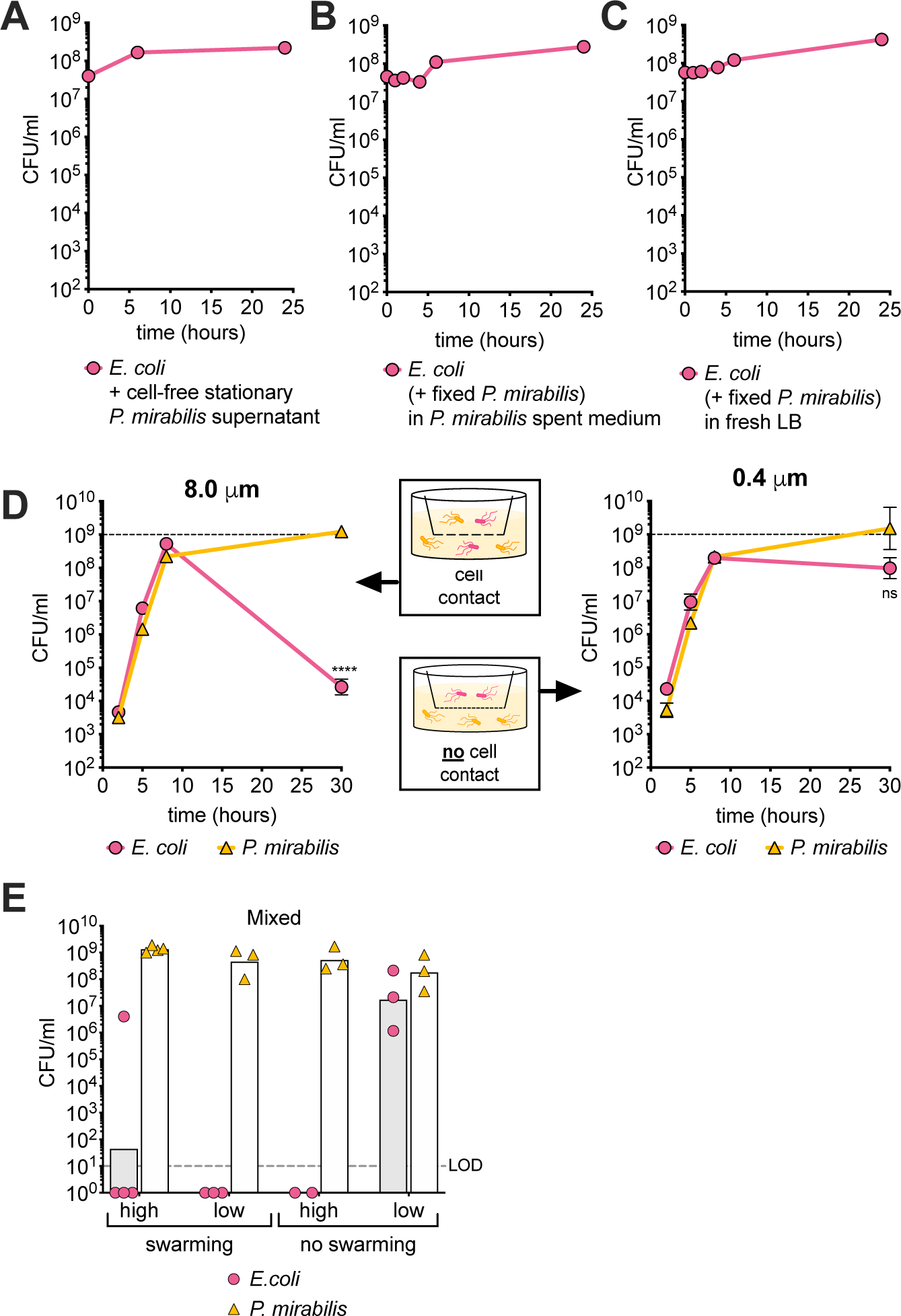
Contact and live cells are required for *P. mirabilis* to kill *E. coli*. **(A-C)** *E. coli* cells grown to stationary phase for 16 hours were added to **(A)** cell-free sterile filtered supernatant of *P. mirabilis*, **(B)** stationary phase *P. mirabilis* supernatant and formalin fixed *P. mirabilis* cells, or **(C)** fresh LB and formalin fixed *P. mirabilis* cells. **(D)** Cultures of *P. mirabilis* and *E. coli* cells in 6-well plates separated by membranes with 8.0 µm or 0.4 µm pore sizes. **(E)** Solid surface assay with mixed cultures of *P. mirabilis* and *E. coli* on swarming permissive (LB) agar or non-swarming permissive (MacConkey) agar in high density (10^8^ CFU of each strain) or low density (10^3^ CFU of each strain) of cells in each spot. All data represent mean of at least three biological replicates (single culture of *E. coli* and high density co-culture on MacConkey two biological replicates). If error bars are not visible, error is smaller than symbol size. For panel D, a Welch’s t-test was performed between the 8-hour time point and the 30-hour time point. ****= p<0.0001, ns = not significant. LOD = limit of detection.

To understand whether killing requires direct contact, we separated *E. coli* and *P. mirabilis* cultures with membranes of different pore sizes (Fig. 2D). An 8.0 µm membrane allows exchange of the secretome and physical contact between cells. We observed about a 20,000 fold reduction in *E. coli* viability at 30 hours compared to 8 hours (Fig. 2D left panel). In contrast, we observed no significant loss in *E. coli* viability with an 0.4 µm membrane that blocked physical contact, only allowing the passage of the secretome (Fig. 2D right panel). *P. mirabilis*-mediated killing is therefore an active process that requires live *P. mirabilis* in direct contact with *E. coli*.

### *P. mirabilis* kills *E. coli* cells on solid surface in a contact-dependent manner

The contact-dependent weapons of Gram-negative species have predominantly been reported to be active and expressed on solid media, with only very few being active in shaking liquid culture (38, 39). We therefore tested *P. mirabilis* and *E. coli* competition on solid media. We inoculated *P. mirabilis* and *E. coli* cells in either high cell density (10^8^ CFU) or low cell density (10^3^ CFU) on two different media. On LB agar, *P. mirabilis* can swarm and cover an entire agar plate within 24h. On MacConkey agar, swarming is inhibited and *P. mirabilis* forms distinct colonies. We recovered similar numbers from all conditions when only one species was present (Fig. S2). When we cultured *P. mirabilis* and *E. coli* together on swarming permissive LB agar, we recovered no viable *E. coli* cells, regardless of the seeding density. On LB agar, *P. mirabilis* cells have the ability to actively move and make physical contact with *E. coli* cells. In contrast, mobility is severely reduced on non-swarming permissive MacConkey agar. Here, *P. mirabilis* eliminated *E. coli* cells in a cell density-dependent manner. When *P. mirabilis* and *E. coli* cells were seeded in high density, and thus with high possibility of bacteria coming in contact with one another, we recovered no viable *E. coli* cells. When cells were seeded in low density and thus physically separated, we recovered significantly more *E. coli* cells (Fig. 2E,F). One possible explanation for this phenotype is that individual cells of *E. coli* and *P. mirabilis* formed microcolonies that eventually touched, but *P. mirabilis* was apparently unable to penetrate the *E. coli* microcolonies and reduce *E. coli* viability.

### Killing is not mediated through the *P. mirabilis* Type VI Secretion System (T6SS)

The T6SS is a well-characterized inter-bacterial killing system (5, 40–42). On solid surfaces *P. mirabilis* uses T6SS against non-kin clonemates (42, 43). However, we observed that *P. mirabilis* killed antagonists in shaking liquid media (Fig. 1). The current literature does not provide evidence that *P. mirabilis* or any other organism uses T6SS in shaking liquid media (13, 43–45). Nevertheless, we tested whether *P. mirabilis* utilizes T6SS to reduce viability of *E. coli*. One key characteristic of *P. mirabilis* is its ability to swarm on permissive agar. T6SS activity will lead to the formation of macroscopically visible zones of dead cells called Dienes lines between strains. Our mouse isolate strain of *P. mirabilis* formed Dienes lines with the type strain for *P. mirabilis* (ATCC #29906) and a human isolate (ATCC #7002) (red arrows, Fig. 3A). The three strains therefore utilize T6SS against non-kin strains. However, during co-culture in shaking liquid media, no reduction in viability of either strain was observed (Fig. 3B,C). To confirm that killing is indeed independent of T6SS, we used a *P. mirabilis* Δ*tssM* mutant, which is deficient in T6SS-mediated effector delivery. This mutant was generated in the *P. mirabilis* strain BB2000, which was shown to encode only a single T6SS (14, 46). Wild type BB2000 and the Δ*tssM* mutant strain reduced the viability of *E. coli* to the same extent and with the same kinetics (Fig. 3D,E). The mechanism used by *P. mirabilis* to reduce viability of *E. coli* is therefore independent of T6SS activity.

**Figure 3:**
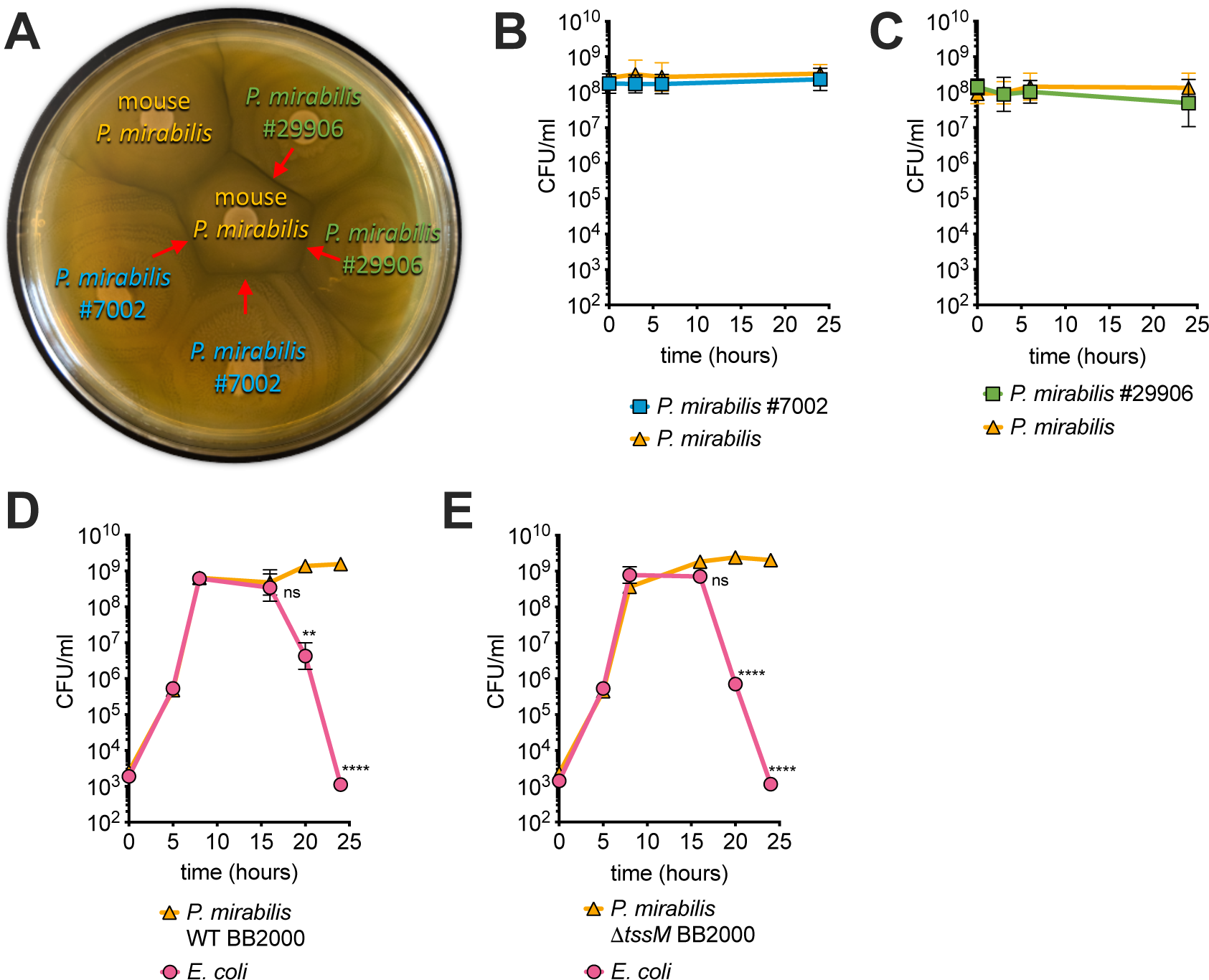
Killing is not mediated through the *P. mirabilis* T6SS. **(A)** Representative image of mouse *P. mirabilis, P. mirabilis* ATCC #7002 and *P. mirabilis* ATCC #29906 that were spotted in duplicate on LB agar and incubated for >24h. Red arrows indicate strong Dienes line formation. **(B+C)** Liquid culture “killing assay” of mouse *P. mirabilis* and **(B)** *P. mirabilis* ATCC #7002 or **(C)** *P. mirabilis* ATCC #29906. **(D+E)** Co-culture in LB of *E. coli* and **(D)** BB2000 WT or **(E)** BB2000 mutant Δ*tssM*, which lacks a functional T6SS. All data represent mean +/-SEM of at least three biological replicates. A one-way ANOVA was performed between the 8-hour time point (upon entry to stationary phase) and the remaining timepoints (stationary phase). **= p<0.01, ****= p<0.0001, ns= not significant

### *P. mirabilis* kills *E. coli* rapidly and without prior contact

*P. mirabilis* does not kill *E. coli* when the co-culture is actively growing in exponential phase. Some contact-dependent killing mechanisms require receptors on recipient surfaces for effector uptake (18, 47), which might not be expressed by *E. coli* during exponential growth phase (48). However, we found that regardless of the growth phase, *E. coli* rapidly lost viability when the cells were added to a stationary phase culture of *P. mirabilis*. After one hour of co-incubation, we recovered about 13,000 and 23,000-fold reduced numbers of stationary phase *E. coli* and exponential phase *E. coli*, respectively (Fig. 4A). *P. mirabilis* therefore kills *E. coli* irrespective of the growth phase of *E. coli*. It also shows that *P. mirabilis* has the ability to kill *E. coli* without prior contact with *E. coli* during exponential phase.

**Figure 4:**
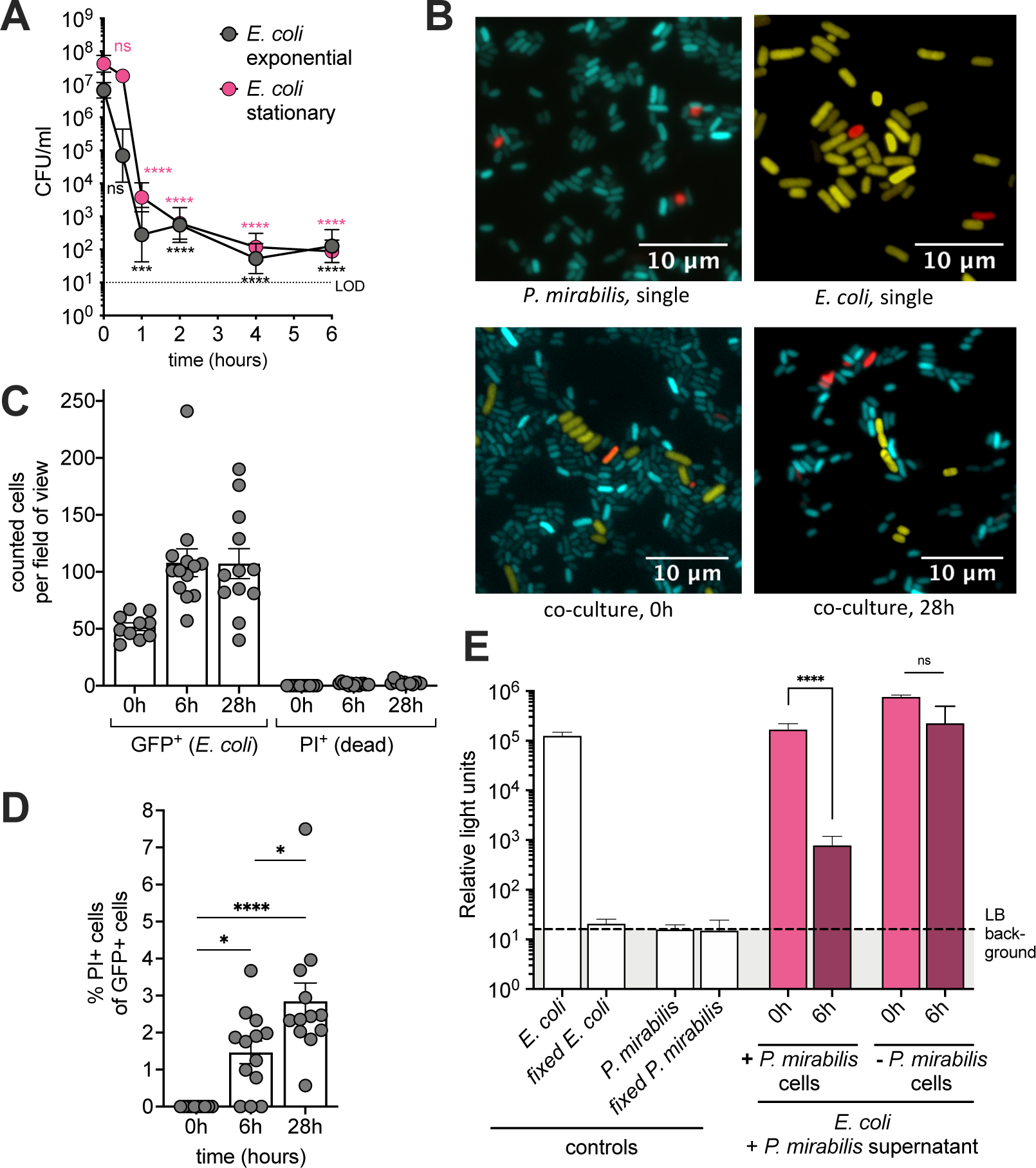
Killing is rapid and does not cause loss of membrane integrity in target cells. **(A)** Exponential phase or stationary phase *E. coli* cells were added to a stationary *P. mirabilis* culture and viability of *E. coli* was determined at indicated time points. A one-way ANOVA was performed comparing the 0-hour time point and remaining time points. **(B)** Representative images of YFP-expressing *E. coli* (yellow) and CFP-expressing *P. mirabilis* (cyan) stained for PI (red), top left to right: monoculture *P. mirabilis,* monoculture *E. coli,* bottom left to right: co-culture at 0 hours and after 28 hours. **(C+D)** GFP-expressing *E. coli* was added to a stationary *P. mirabilis* culture (= “killing assay”) and culture was imaged 0 hours, 6 hours or 28 hours after beginning of co-culture: **(C)** Number of GFP-positive (*E. coli*) or propidium iodide (PI) positive (*P. mirabilis* and *E. coli*) cells per field of view. **(D)** Percentage of GFP-positive (*E. coli*) cells that stain positive for PI. **(E)** Bioluminescent *E. coli* LuxAB in stationary phase was added to a stationary phase *P. mirabilis* and bioluminescence measured immediately and after 6 hours. Controls: single culture *E. coli* LuxAB and single culture *P. mirabilis,* untreated or fixed in formalin. All data represent mean +/-SEM of at least three biological replicates. For panel B, an ordinary one-way ANOVA test and for panels C and D Tukey’s multiple comparison test was used. *=p<0.05, **= p<0.01, ***= p< 0.001,****= p<0.0001, ns= not significant.

### *E. coli* cells maintain cell shape and integrity despite loss of viability and metabolic activity

To gain a better understanding of the potential killing mechanism used by *P. mirabilis*, we used fluorescence microscopy. We imaged a co-culture of Cyan Fluorescent Protein (CFP)-expressing *P. mirabilis* and Yellow Fluorescent Protein (YFP)-expressing *E. coli* and used propidium iodide (PI) to test for membrane integrity of cells. After 28 hours of co-incubation, we did not observe changes in the number of YFP-expressing cells, their cellular morphology, or an increase of PI-positive cells compared to single cultures (Fig. 4B). We also quantified our findings using GFP-expressing *E. coli*. The number of *E. coli* cells that we observed in each field of view by fluorescence microscopy was not reduced (Fig. 4C).

According to CFU counts (Fig. 1G, 4A), at 6h of co-culture almost all *E. coli* cells in each field of view should be non-viable. However, we observed very little PI uptake (1.5%) of GFP-positive (*E. coli*) cells (Fig. 4C+D). After 28h of co-culture, the percentage of PI-positive and GFP-positive cells increased only slightly to 2.9% (Fig. 4C+D). The cellular morphology of GFP-positive cells also remained unchanged throughout the experiment (Fig. S3). We next investigated if these *E. coli* cells were still metabolically active using bacterial bioluminescence. When cells stop producing ATP, bioluminescence is rapidly lost. When bioluminescent *E. coli* cells were added to stationary phase culture of *P. mirabilis,* we observed a significant reduction in bioluminescence (Fig. 4E). No significant reduction in bioluminescence occurred in stationary phase *P. mirabilis* supernatant in the absence of *P. mirabilis* cells (Fig. 4E). Taken together, these data suggest that *P. mirabilis* compromises metabolic activity of *E. coli,* but the mechanism does not result in a compromised cellular envelope in target cells.

### Activity of the killing system is differentially regulated in stationary phase

*P. mirabilis* rapidly reduces viability of *E. coli* in a stationary phase co-culture (Fig. 4A). Interestingly, the number of remaining viable *E. coli* cells remains largely unchanged between 20-30 hours post inoculation (Fig 1B). These remaining *E. coli* cells did not acquire resistance, as they were killed when isolated, grown, and exposed again to *P. mirabilis* (data not shown). One possible explanation for the sustained viability of *E. coli* cells is that *P. mirabilis* downregulates expression of the killing system. Indeed, the capability of *P. mirabilis* to reduce viability of added *E. coli* gradually diminished during stationary phase (Fig. 5A). When *P. mirabilis* cells were grown for 25 hours, they lost the ability to kill *E. coli* cells (Fig 5A). At this time point, a culture of *P. mirabilis* also did not reduce the viability of *S*. Typhimurium (Fig. S4). Taken together, these data suggest that as a *P. mirabilis* culture ages, cells remain viable but lose the ability to kill *E. coli*.

**Figure 5:**
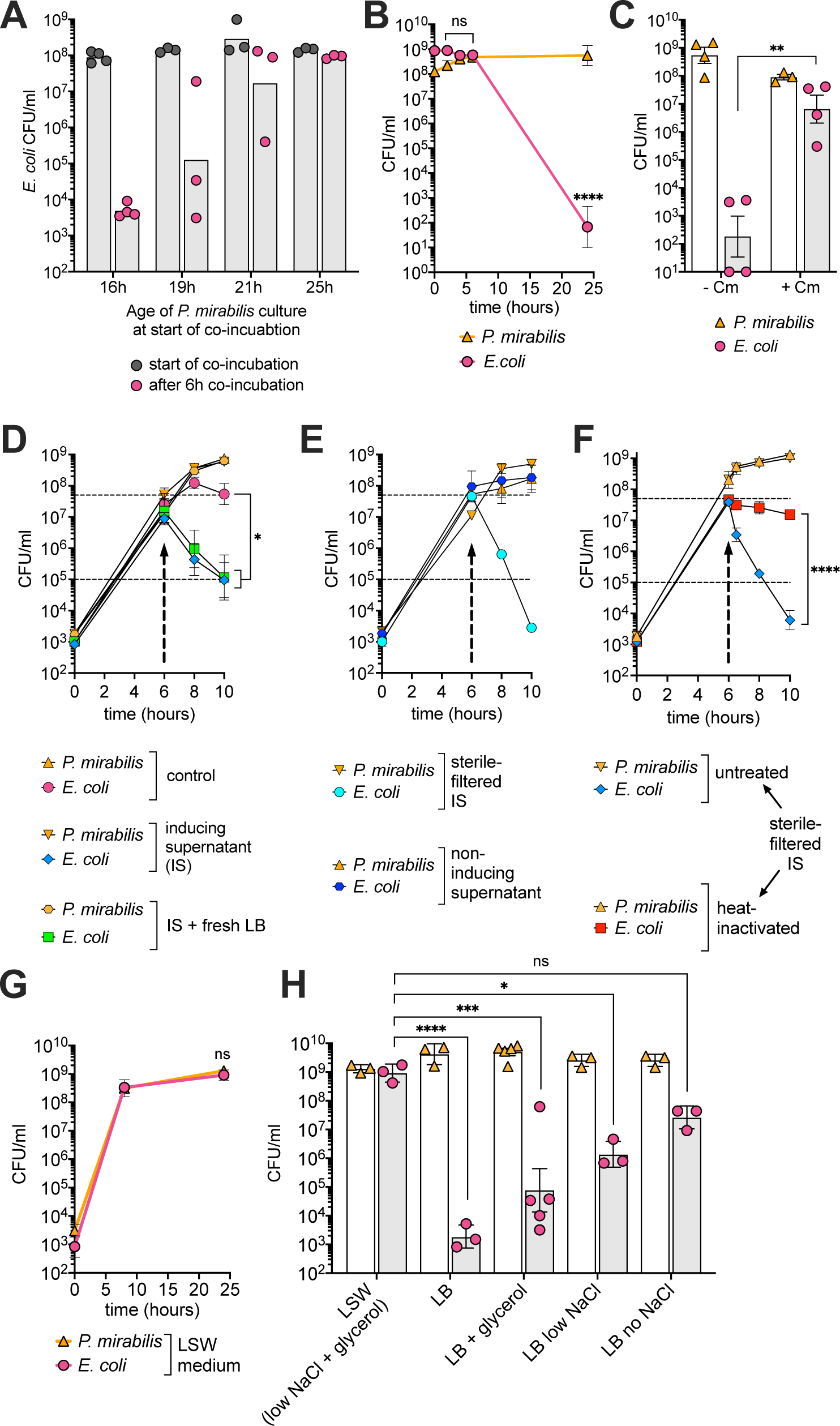
Killing requires new protein synthesis and is regulated by a component in the culture supernatant and osmolarity. **(A)** *P. mirabilis* was grown in single culture for 16, 19, 21, or 25 hours before stationary phase *E. coli* cells were added. Viability of *E. coli* was assessed immediately or 6 hours after start of co-incubation. **(B)** *E. coli* was grown in single culture for 16 hours before stationary phase *P. mirabilis* cells were added. **(C)** *E. coli* was grown in single culture for 16 hours before stationary phase *P. mirabilis* cells and chloramphenicol (15 µg/ml) were added. Histogram bar represents viability 24 hours after the beginning of co-culture. **(D-F)** Killing induction assays. All data represent mean +/-SEM of at least three biological replicates. For panels C and F, a Welch’s t-test was performed. For panel D, a one-way ANOVA was performed comparing viability of *E. coli* at the time of supernatant exchange and remaining time points. Upward arrow marks time of supernatant exchange. The medium of a co-culture of *P. mirabilis* and *E. coli* grown for 6 hours (indicated by arrows) was replaced with: **(D)** The growing culture’s own supernatant (control). Supernatant of 22-hour old culture of *P. mirabilis* (inducing supernatant, IS). 1:1 mixture of fresh LB and inducing supernatant (IS + fresh LB). **(E)** Sterile-filtered supernatant of 22-hour old culture of *P. mirabilis* (sterile filtered IS). Supernatant of 22-hour old culture of *E. coli* (non-inducing supernatant). (**F**) Sterile-filtered supernatant of 22-hour old culture of *P. mirabilis* (sterile filtered IS) either not treated (untreated) or boiled for 15 min (heat-inactivated). **(G)** Shaking liquid growth curve of mouse *P. mirabilis* and *E. coli* in LSW (low NaCl, glycerol) broth. A Welch’s t-test was performed between the 8-hour time point and the 24-hour time point comparing viability of *E. coli*. ns= not significant. **(H)** Viability of *E. coli* and *P. mirabilis* after 24 hours of co-culture in different media. A one-way ANOVA was performed comparing the viability of *E. coli* in different media to viability in LSW. *= p<0.05, **= p<0.01, ***= p<0.001 ****= p<0.0001, ns= not significant.

The viability of *E. coli* cells was quickly reduced when placed in a stationary phase *P. mirabilis* culture (Fig 1G, 3A, 4A and 4B). However, when the inoculation was reversed and *P. mirabilis* cells were placed in a stationary culture of *E. coli*, killing was delayed. In this reversed setting, *E. coli* cells were alive 6 hours post incubation with *P. mirabilis* cells (Fig 5B). This prompted us to test whether new protein synthesis is required for killing. We used chloramphenicol to inhibit new protein synthesis. A chloramphenicol resistant strain of *E. coli* was grown for 16 hours, before 15µg/ml of chloramphenicol and the chloramphenicol sensitive *P. mirabilis* were added. This concentration of chloramphenicol is above the minimum inhibitory concentration (7.8µg/ml) but does not interfere with *P. mirabilis* viability (Fig. 5C). Addition of chloramphenicol resulted in a significant (about 40,000-fold) rescue in viability of *E. coli* compared to the control group without chloramphenicol (Fig. 5C), indicating that new protein synthesis is required for killing.

### Component(s) of *P. mirabilis* supernatant regulate expression of the killing system

*P. mirabilis* cells need to reach a high cell density before killing is observed (Fig. 1B and 5B). This might indicate that the behavior is regulated by quorum sensing (QS). QS is a form of bacterial cell-to-cell communication that involves the production, release and response to extracellular molecules that collectively alter bacterial behavior (49). *P. mirabilis* cells might secrete QS molecules that accumulate in the supernatant as *P. mirabilis* culture density increases. These molecules might regulate expression, post-translational modification or use of the killing proteins. To test this hypothesis, we grew *P. mirabilis* and *E. coli* in a co-culture to mid-exponential phase (6 hours). At this point, we replaced the growing culture supernatant with the supernatant of a stationary phase (22-hour) culture of *P. mirabilis*, which could harbor putative QS molecules. This change indeed resulted in loss of viability of *E. coli* in the co-culture. Compared to the control group, we observed a highly significant 467-fold reduction in the viability of *E. coli* cells 4 hours after the supernatant exchange (Fig. 5D blue line vs pink line). Nutrient deprivation and starvation were not responsible for the observed phenotype, as killing was also induced to the same level with a 50:50 mixture of *P. mirabilis* stationary culture supernatant and fresh LB (Fig. 5D green line). *E. coli* supernatant (non-inducing) did not result in loss of viability of *E. coli* (Fig. 5E). Sterile-filtered supernatant of *P. mirabilis* resulted in the same, if not even greater, viability loss of *E. coli* as non-sterile filtered supernatant (Fig. 5D,E). We noticed that the proposed signaling molecule in the supernatant seems to be heat labile, as boiling of the 22-hour *P. mirabilis* supernatant was able to partially rescue *E. coli* viability (Fig. 5F red line). Although *P. mirabilis* supernatant alone was not sufficient in killing *E. coli* (Fig. 2A), there seems to be a heat labile component within the supernatant that accumulates during the transition to stationary phase that regulates the *P. mirabilis* killing mechanism.

### The killing system is regulated via osmolarity

The killing system also seems to be regulated by additional environmental factors. We discovered one such factor when we used the common *P. mirabilis* medium L swarm minus (LSW) (50) that inhibits swarming. When we performed solid surface killing on LSW agar, *P. mirabilis* failed to kill *E. coli* (not shown). In a co-culture in LSW broth *P. mirabilis* also failed to kill *E. coli* (Fig. 5G). LSW and LB broth differ not only in their NaCl concentrations, but also in glycerol as an additional carbon source in LSW. We therefore investigated the contribution of the individual components. In LB media supplemented with glycerol alone, *E. coli* was killed to almost the same extent as in LB. However, in LB with reduced NaCl concentration (0.4g), we observed a significant increase in the viability of *E. coli* cells. Complete omittance of NaCl from LB resulted in an even greater survival of *E. coli* cells. However, there was a trend of lower survival in LB without NaCl compared to LSW (Fig. 5H), suggesting that both low NaCl concentration and the presence of glycerol act to inhibit the expression and/or use of the killing system.

## DISCUSSION

### Novelty of the killing system

Although a number of bacterial competition systems have been discovered, our understanding of bacterial warfare is still limited (12). The majority of contact dependent killing mechanisms have been discovered and studied over just the past 15 years and a vast majority of organisms could harbor potentially novel mechanisms of competition (12). We initially observed what was apparently competition between a commensal *P. mirabilis* strain and a commensal *E. coli* in neonatal mice. During studies *in vitro*, we replicated the *in vivo* observation and showed that *P. mirabilis* killed competing *E. coli*. We have discovered that *P. mirabilis* utilizes a contact-dependent system to kill prey species (Fig 2A,D). The known contact dependent arsenal of weapons used by Gram-negative bacterial species against other species include T3SS, T4SS, T6SS, CDI (T5SS), Cdz, and MccPDI (12, 22). The inter-species killing system we describe is novel for *P. mirabilis*, as it is independent of T6SS (Fig. 3E). The system might also represent a novel system for bacteria in general or an extension of a known system, as the characteristics significantly differ from known contact dependent mechanisms. We outline these differences in the following.

*P. mirabilis* encodes a T3SS (51). T3SS are widely used by bacteria to communicate with organisms belonging to other kingdoms, including protists, fungi, plants and animals (52). While some bacteria use T3SS to kill eukaryotic yeast cells (53, 54), current literature does not provide evidence for the use of T3SS as an interbacterial competition system. Alternatively, the T4SS has been described as an inter-species killing system for *Stenotrophomonas* and *Bartonella*, but it was only effective on solid media (55, 56).

The T6SS is widely used by Gram-negative species as an intra- and inter-bacterial killing mechanism (5, 40, 43, 46, 57–61) and represents the only contact-dependent killing mechanism described for *P. mirabilis* (51). However, activity of *P. mirabilis* T6SS has only been shown against non-kin clonemates and not against other species (42, 62). As our study demonstrates, a mutant strain of *P. mirabilis* unable to produce a functional T6SS was able to kill *E. coli* (Fig. 3E)*. P. mirabilis* therefore utilizes a mechanism that differs from the canonical T6SS.

Contact dependent growth inhibition (CDI) depends on a two-partner secretion system and was first identified in *E. coli* (CdiA/CdiB) (38). The CDI system arrests growth in the target species but does not reduce viability. Moreover, the attacking strain has to be in exponential phase to deliver a toxic protein to the target cell. When the attacking strain is in stationary phase, as we see for *P. mirabilis*, it does not inhibit the target strain (38). Contrary to our data showing that a series of *Enterobacteriaceae* species are killed by *P. mirabilis* (Fig. 1), the effects of *E. coli* CDI do not expand to non-related species (18, 63, 64). Furthermore, CDI has been described to be predominantly active on solid media and not in shaking liquid culture (17, 20). One study on *P. aeruginosa* CDI showed some activity in liquid media. However, inhibition of susceptible strains is significantly reduced compared to solid surface conditions (39). It seems unlikely that *P. mirabilis* uses a CDI-like system against *E. coli*, as A) killing of *E. coli* did not occur in exponential phase, B) distantly related species are killed, C) contact dependent killing occurred both on solid surface and in shaking liquid media.

Another mechanism used by Gram-negative bacteria is contact-dependent inhibition by glycine zipper proteins (Cdz) (21). *Caulobacter crescentus* utilizes Cdz against susceptible *C. crescentus* strains and a limited number of species in the *Caulobacteraceae* (47). The catalytic bacteriocin-like proteins CdzC and CdzD transfer through a receptor, presumably PerA (47) into the recipient cells and cause rapid depolarization of the cell membrane. An active Cdz system caused the vast majority (>95%) of target cells to lose membrane integrity, round up, and stain positive for propidium iodide (21). We did not observe a considerable uptake of PI or any changes in *E. coli* cellular morphology. Thus far, only the effector proteins CdzC and CdzD have been described for the Cdz system. *P. mirabilis* might use other effector proteins that differ in their mechanism of action against target bacteria. However, this is unlikely, as the authors of the study did not find homologs of the Cdz system in *Proteus* spp. (21).

Microcin proximity-dependent growth inhibition (MccPDI), identified in *E. coli* (22), occurs in shaking liquid media and requires direct contact or proximity between cells. However, killing or inhibition by the attacking strain occurs in late exponential phase. Moreover, killing seems to be limited to different *E. coli* isolates (65). Finally, MccPDI requires the outer membrane protein OmpF on the target cells (66). Deletion of *ompF* rendered strains resistant and expression of compatible OmpF rendered *Salmonella enterica* and *Yersinia enterocolitica* susceptible to *E. coli* MccPDI (66). However, *P. mirabilis* also killed an OmpF deficient strain of *E. coli* (data not shown). Thus, the mechanism used by *P. mirabilis* to inhibit *E. coli* seems to be independent of MccPDI.

### Regulation of the killing system

*P. mirabilis* reduced viability of target species shortly after entering stationary phase. Presumably, the expression of the killing system itself or its regulator is upregulated upon reaching a concentration of above 10^9^ CFU/ml (Fig. 1). Interestingly, we found that supernatant of *P. mirabilis* in stationary phase harbors component(s) that might function in a regulatory circuit controlling the activity of the killing system. Stationary phase *P. mirabilis* culture supernatant was sufficient to induce killing by *P. mirabilis* in a co-culture still in exponential phase. The component(s) in the supernatant might be signaling molecules in a quorum sensing (QS) regulatory network. As bacteria grow, they release QS molecules into their extracellular environment. Upon accumulation of these extracellular molecules, bacteria collectively alter many of their physiological behaviors such as competence, biofilm formation, and secretion systems (49, 67). Gram-negative microbes such as *V. cholerae, P. aeruginosa, E. coli,* and *Serratia liquefaciens* regulate their secretion systems, especially those known to be involved in bacterial killing, through QS (68). Many species of *Pseudomonas* and *Vibrio* regulate their killing mechanisms through QS molecules (58, 69–74). Gram-negative species have a plethora of mechanisms to synthesize QS molecules that include production of homoserine lactones (HSL) by LuxI/RpaI and autoinducer-2 (AI-2) by LuxS genes (49). *P. mirabilis* (strain BB2000) is known to harbor two genes involved in QS. The first is LuxS, encoding for AI-2 molecules, and the second is a receptor, RbsA, a homolog of LuxQ in *Vibrio* species (75). However, very few targets for QS systems in *P. mirabilis* have been identified. LuxS was reported to initiate swarming of *P. mirabilis in vitro*. A second report found a role for exogenously added HSLs in biofilm formation in *P. mirabilis* (76). However, there are no homologs of any genes encoding for HSLs in the genome of *P. mirabilis*. Nonetheless, these reports suggest a role for AI-2s and HSLs in regulating QS pathways in *P. mirabilis*. AI-2s or HSLs could therefore potentially be involved in the regulation of the *P. mirabilis* killing mechanism. The killing system also appears to be regulated by environmental factors such as osmolarity and carbon source (Fig. 5H). Further studies are required to determine the nature and the mechanism by which putative signaling molecules and environmental factors govern the killing system used by *P. mirabilis*.

### Biological relevance

Our study focuses on the *in vitro* characterization of how *P. mirabilis* reduces viability of competitor species. However, it was informed by an *in vivo* observation in the mouse model with mouse-adapted gut commensal species. Our *in vitro* findings indicate that the system is active when *P. mirabilis* reaches a high cell density. In infants of mice and humans, *Enterobacteriaceae* species like *P. mirabilis* and *E. coli* initially dominate the microbiota and reach densities similar to stationary phase growth *in vitro* (10^9^ CFU/g of fecal matter) (77, 78). It thus seems probable that *P. mirabilis* expresses the contact-dependent killing system in the infant gut and kills the competitor *E. coli*. However, future studies are needed to determine under which conditions *P. mirabilis* uses this killing system in the gut.

The two species used in our study were isolated as commensals from the gastrointestinal tract of mice. However, both *P. mirabilis* and *E. coli* are also well-known causes of urinary tract infections of humans. *P. mirabilis* is a concern for catheter-associated urinary tract infections (CAUTI) and is frequently found on the catheter surface during long-term catheterization (79, 80). Polymicrobial biofilms are rarely seen during short-term catheterization, but their incidence increases to up to 77% during long-term catheterization (51, 79). One study found *E. coli* to be only rarely (18 %) associated with *P. mirabilis* on catheters (81), while another study found *P. mirabilis* more frequently associated with *E. coli* (38 % of specimens, 45% of patients) (82). Whether *P. mirabilis* employs its putative inter-species competition system also in mixed biofilms will be an interesting question to address in future studies.

### Concluding remarks

We report that *P. mirabilis* is able to reduce viability of a variety of *Enterobacteriaceae* species in a contact-dependent manner. The reduction of viability is independent of the T6SS, the only contact-dependent killing system described for *P. mirabili*s. To our knowledge, the mechanism is novel for *P. mirabilis*. The present study is a first description and characterization of the observed *P. mirabilis*-mediated inter-bacterial killing phenotype. The identity of the genetic components required for the expression of the system are still unknown and the focus of ongoing studies. Similarly, the current study indicates possible regulatory mechanisms for the system, which are being investigated. As four different *P. mirabilis* isolates reduced viability of *E. coli*, it might be a capability that is widespread among *P. mirabilis* strains. Frequently, competition systems are also not restricted to one species but can be found in multiple species, as shown for, e.g., T6SS, T4SS, CDI, and Cdz. The competition system encoded by *P. mirabilis* might therefore also be present in other species.

## MATERIALS AND METHODS

### Mouse and bacterial strains

The female C57BL/6 SPF mouse used in the breeding originated from a rederivation with Envigo CD-1 mice and thus harbors CD-1 microbiota. The male C57BL/6 SPF mouse used in the breeding was obtained from Taconic laboratories. Both the *Escherichia coli* and *Proteus mirabilis* isolated from these mice are natural colonizers of the GI tract of these animals. The dam and the sire were allowed to mate, and both were kept in the same cage before the pups were weaned at approximately 21 days of age. Mouse breeding and isolation of bacterial strains were performed at the University of California, Irvine. All animal experiments were reviewed and approved by the Institutional Animal Care and Use Committee at the University of California, Irvine.

**Table 1:**
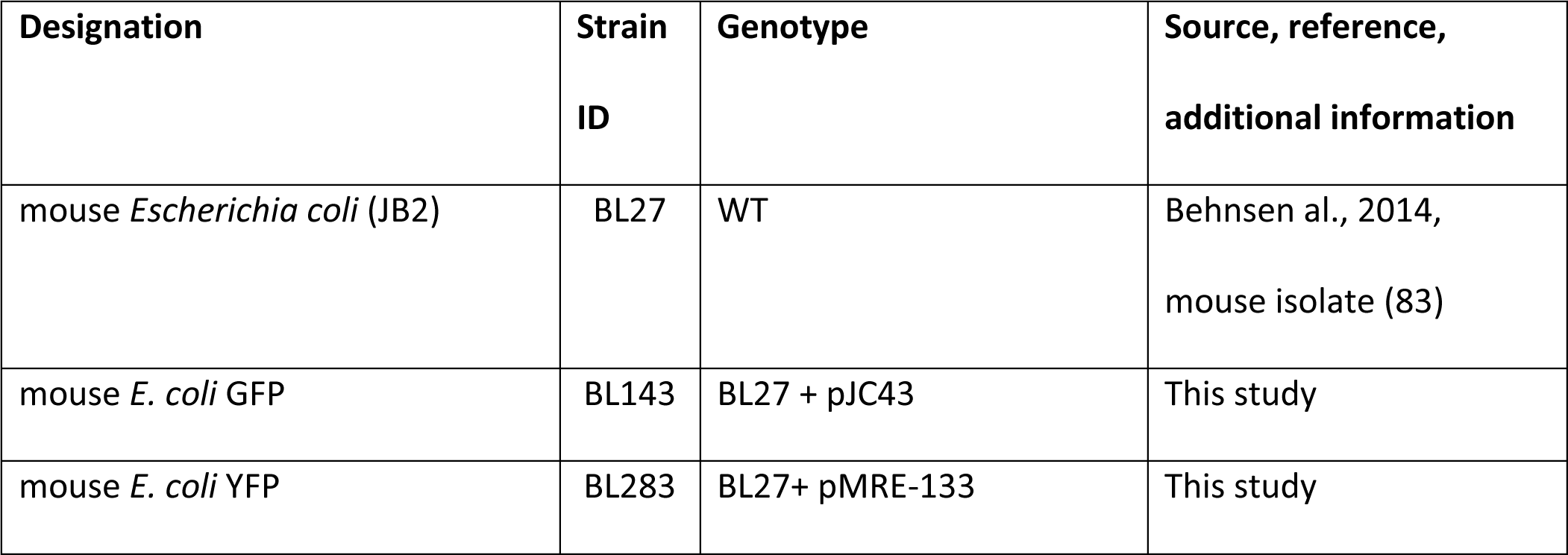
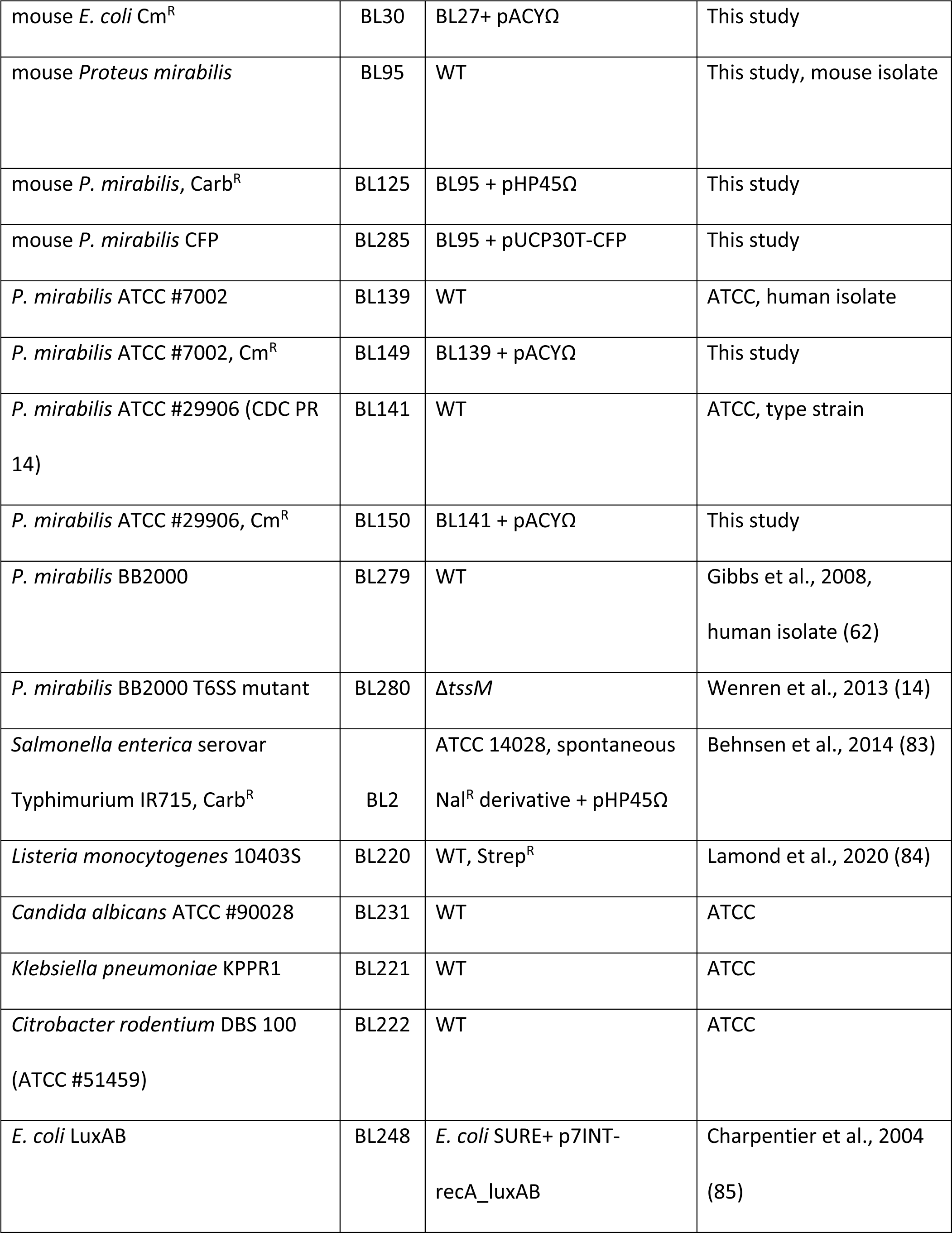
Bacterial strains

**Table 2:**
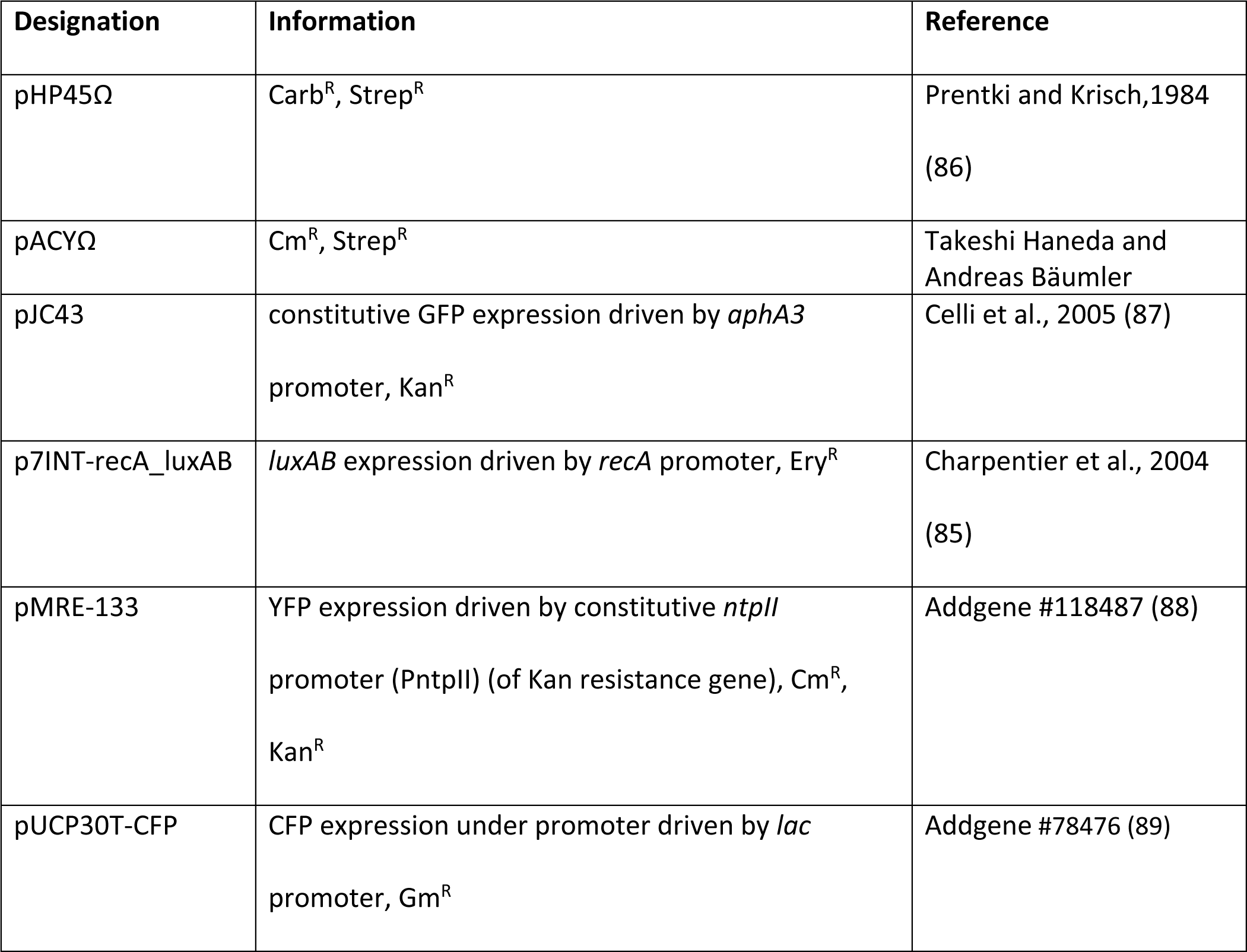
Plasmids

### Media and growth conditions

*P. mirabilis, E. coli, C. rodentium*, *S.* Typhimurium, and *K. pneumoniae* were routinely cultured in Lysogeny Broth (LB). *C. albicans* was cultured in Yeast Peptone Dextrose (YPD) broth. *L. monocytogenes* was cultured in Brain Heart Infusion (BHI) broth. Cultures were grown shaking in a 20 mm diameter culture tubes at 200 rpm for 16 hours in 5 ml of medium unless otherwise indicated. *P. mirabilis* BL125 was selected on MacConkey agar plates supplemented with carbenicillin (100 mg/L). *P. mirabilis* strains BL149 and BL150 were selected on MacConkey agar supplemented with chloramphenicol (30 mg/L). *E. coli* strains BL143 and BL30 were selected on MacConkey agar plates supplemented with kanamycin (50 mg/L) and chloramphenicol (30 mg/L) respectively. *S*. Typhimurium was selected on LB agar supplemented with carbenicillin (100 mg/L). *K. pneumoniae* cells were selected on MacConkey agar supplemented with carbenicillin (100 mg/L). *L. monocytogenes* was selected on BHI agar plates supplemented with streptomycin (100 mg/L). *P. mirabilis* strains BL95, BL279 and BL280 were selected for on MacConkey agar plates with no antibiotics. *C. albicans* was selected for on Sabouraud agar supplemented with carbenicillin (100 mg/L). For the LSW co-cultures, BL95 was selected for on LB agar plates with tetracycline (20 mg/L).

### Monoculture growth assays

*E. coli* BL143 was grown in LB for 16 hours. The strain was centrifuged at 9400 rcf, resuspended, and diluted in sterile PBS. Twenty ml LB were inoculated with 5 x 10^3^ CFU/ml of BL143. The culture was incubated shaking at 200 rpm at 37°C, and an aliquot (100 µl) was collected at 0, 2, 5, 8, 16 and 24 hours. This aliquot was resuspended in sterile PBS, serially diluted, and plated on MacConkey agar.

### Co-culture growth assays

*P. mirabilis* BL95 and its respective antagonist were grown separately in LB for 16 hours. Each strain was centrifuged at 9400 rcf, resuspended, and diluted in sterile PBS. Twenty ml LB were inoculated with 5 x 10^3^ CFU/ml of *P. mirabilis* BL95 and antagonist. The co-culture was incubated shaking at 200 rpm at 37°C, and an aliquot (100 µl) was collected at 0, 2, 5, 8, 10, 14, 16, 18, 20, 24 and 30 hours. This aliquot was resuspended in sterile PBS, serially diluted, and plated on agar plates with antibiotics to select for *P. mirabilis* or the antagonist.

### Killing assay

*P. mirabilis* and Gram-negative antagonist species were grown separately in 5 ml of LB for 16 hours shaking at 37°C. *L. monocytogenes* was grown in 5ml of Brain Heart Infusion (BHI) at 37°C, and *C. albicans* was grown in 5 ml of Yeast Peptone Dextrose (YPD) at 30°C in shaking liquid culture. 5 x 10^8^ cells of the antagonist were centrifuged (9400 rcf) and resuspended in 500 µl of sterile PBS. The cells were added to 4.5 ml of the *P. mirabilis* culture. This roughly corresponds to an attacking to target ratio of 10:1. The viability of the antagonists was measured by plating on their respective media supplemented with antibiotics (see section above).

### Supernatant assay

*P. mirabilis* BL95 and *E. coli* BL143 cells were grown for 16 hours. *P. mirabilis* culture was centrifuged at 3220 rcf for 15 minutes and the supernatant was sterile filtered using a 0.22 µm syringe filter. 10^8^ *E. coli* cells were centrifuged at 9400 rcf for 5 minutes and resuspended in PBS. *E. coli* cells were added to the sterile filtered supernatant of *P. mirabilis.* The viability of *E. coli* cells was measured 6- and 24-hours after start of incubation in *P. mirabilis* supernatant.

### Split well assay

In this assay, cultures were separated by membranes of different pore sizes in 6-well plates (VWR). Inserts with membrane pore size of 0.4 µm (VWR 10769-192) or 8.0 µm (VWR 10769-196) were placed in wells of the plate, creating two chambers. 2 ml of fresh LB was added to each chamber. Cultures (16h-old) of BL30 and *P. mirabilis* cells BL125 were centrifuged (9400 rcf), resuspended, and diluted in sterile PBS. The upper and lower chamber were inoculated with 5 x 10^3^ cells of *E. coli* BL30 and *P. mirabilis* BL125, respectively. Plates were incubated shaking at 90 rpm at 37°C. The viability of the cells in each chamber was measured by plating the cells on medium selective for either *P. mirabilis* or *E. coli*.

### Solid surface assay

*P. mirabilis* BL95 and *E. coli* BL143 cells were grown in shaking liquid culture for 16 hours. 10 µl of each strain containing either 10^8^ (high density) or 10^3^ (low density) cells were mixed and immediately spotted either on MacConkey agar or LB agar. The plates were incubated at 30°C for 24 hours. After 24 hours, a 9 mm diameter hole puncher was used to stab the center of the growing colony. The agar plug was placed in 2 ml of PBS and cells were resuspended by gentle vortexing. Cells were serially diluted and plated on medium selective for either *P. mirabilis* or *E. coli*.

### Formalin treatment

*P. mirabilis* BL95 was grown for 16 hours and 5 ml of the culture was centrifuged (9400 rcf for 10 minutes). The supernatant was sterile filtered. The cells were resuspended in 1 ml of 10 % formalin, incubated for 5 minutes at room temperature, and washed twice with 1 ml PBS to remove any residual formalin. After the final wash, cells were either resuspended in 5 ml LB or the original sterile filtered *P. mirabilis* culture supernatant. *E. coli* was grown for 16 hours and 5 x 10^8^ cells were added to each tube. The cultures were placed in a shaker at 37°C shaking at 200 rpm and viability of *E. coli* cells was measured by plating serial dilutions on selective media at 6 and 24 hours after the start of co-culture.

### Chloramphenicol treatment

*P. mirabilis* BL95 and *E. coli* BL30 cells were grown for 16 hours. 5 x 10^8^ *P. mirabilis* cells were centrifuged (9400 rcf for 5 minutes) and resuspended in sterile PBS. Five hundred microliters of *P. mirabilis* cells were placed into 4.5 ml of an *E. coli* culture grown for 16 hours. Chloramphenicol (15 µg/ml) was then added to the co-culture. Samples were taken over a 24-hour period, serially diluted and plated on selective media to assess viability. *E. coli* and *P. mirabilis* were selected on MacConkey agar with chloramphenicol and MacConkey with no antibiotics respectively.

### Fluorescence microscopy

*E. coli* BL143 cells expressing green fluorescence protein (GFP) and *P. mirabilis* BL95 were inoculated in a “killing assay” (see respective section). At indicated time points, 100 µl of the co-culture was centrifuged at room temperature for 10 minutes at 9400 rcf. The pellet was resuspended in 1 ml of propidium iodide (PI) in sterile PBS (final concentration of 2 x 10^-3^ mg/ml) and incubated for 15 minutes protected from light. Cells (10 µl) were spotted on a 1x1 cm wide, 5 mm thick agarose pad (1.5 % agarose in PBS) positioned on a microscope slide and incubated 10 minutes to dry. Once the sample was absorbed onto the agarose pad, a cover slip was added and sealed with nail polish. Control cells were treated with 95 % EtOH before addition of PI. The cells were visualized at 60x in a Keyence inverted microscope with GFP (excitation 470/40 emission 525/50 nm) and Texas Red (excitation 560/40 emission 630/75 nm) filter sets. 10 representative images were taken per slide, averaging 100-300 GFP-positive cells per field. Each image was analyzed for the total number of GFP positive and PI positive cells. *E. coli* cells expressing YFP (BL283) were grown for 16 hours with 0.5 % arabinose. *P. mirabilis* cells carrying CFP (BL285) were grown for 16 hours with 1 mM IPTG. A “killing assay” as described above was performed. Representative images of the assay were taken at the start of the assay (0 hour) and 28 hours later. A representative image of the single culture of each strain was taken 28 hours after inoculation. Images were taken with Zeiss AxioImager Z2 upright microscope. The cells were visualized at 63x/1.40 oil DIC M27 magnification with CFP (excitation 433, emission 475 nm) and YFP (excitation 508, emission 524 nm) and RFP (excitation 590, emission 612 nm) filter sets.

### Luminescence assays

Luminescent *E. coli* BL248, non-luminescent *E. coli* BL27, and *P. mirabilis* BL95 were grown for 16 hours. The *P. mirabilis* culture was centrifuged at 3220 rcf for 15 minutes. 4.5 ml of the supernatant was sterile filtered using 0.2 µm syringe filter (VWR 28145-501). 5 x 10^8^ cells of luminescent *E. coli* BL248 were centrifuged (5 minutes 9400 rcf), resuspended in 500 µl of sterile PBS and inoculated either into the sterile filtered supernatant of *P. mirabilis* or 4.5 ml of *P. mirabilis* (“killing assay”). 1 x 10^9^ *E. coli* BL248 and *P. mirabilis* were centrifuged (5 minutes at 9400 rcf), resuspended in 500 µl of formalin and incubated 5 minutes. Formalin treated cells were washed once with 500 µl of sterile PBS. 100 µl (two technical replicates) of each condition were aliquoted into 96 well plates (Greiner bio-one 655083) and exposed to decyl aldehyde (Sigma A0398589) fumes. Luminescence was measured using a Biotek Synergy 2 with the following settings: Integration time: 0:01.00 (MM:SS.ss), Filter Set 1 Emission: Hole Optics:Top Gain: AutoScale, Read Speed: Normal, Delay 100msec. The average between each technical replicate was designated as one biological replicate.

### Killing induction assays

To obtain supernatants for killing induction, a culture of *P. mirabilis* BL95 was grown for 22 hours. Cells were centrifuged at 3220 rcf for 20 minutes and supernatants were obtained by decanting. Supernatants were either used directly in assays or were sterile filtered before use. For the killing induction assay, *E. coli* cells (BL143) and *P. mirabilis* cells (BL95) were grown for 16 hours separately. 10^10^ cells of each strain were centrifuged at 9400 rcf for 5 minutes and resuspended in 500 µl of sterile PBS. 5 x 10^3^ cells of each strain were inoculated in 5 ml of fresh LB and incubated shaking at 37°C and 200 rpm. The co-culture was grown for 6 hours and centrifuged at 3220 rcf for 15 minutes. The pellet was resuspended in 5 ml of different media: its own supernatant, 22-hour supernatant of *P. mirabilis* (inducing supernatant), 2.5 ml of fresh LB and 2.5 ml of inducing supernatant, boiled inducing supernatant (95°C for 15 minutes), 22-hour supernatant of *E. coli*, or 22-hour sterile-filtered supernatant of *P. mirabilis*.

### Co-culture assay in LSW

*P. mirabilis* BL95 and *E. coli* BL143 were grown in LSW for 16 hours. Each strain was centrifuged at 9400 rcf, resuspended, and diluted in sterile PBS. Twenty ml LSW was inoculated with 5 x 10^3^ CFU/ml of each strain. The co-culture was incubated shaking at 200 rpm at 37°C, and an aliquot (100 µl) was collected at 0, 8 and 24 hours. This aliquot was resuspended in sterile PBS, serially diluted, and plated on agar plates with antibiotics to select for *P. mirabilis* (LB + tetracycline) or *E. coli* (MacConkey + kanamycin). To investigate the individual components of LSW broth, the following media were prepared, the pH adjusted to 7, and autoclaved.

**Table 3:** Media composition

|  | NaCl (g/L) | Tryptone (g/L) | Yeast Extract (g/L) | Glycerol (ml/L) |
| --- | --- | --- | --- | --- |
| LB | 10g | 10g | 5g | 0ml |
| LSW | 0.4g | 10g | 5g | 5ml |
| LB glycerol | 10g | 10g | 5g | 5ml |
| LB low NaCl | 0.4g | 10g | 5g | 0ml |
| LB no NaCl | 0g | 10g | 5g | 0ml |

### Swarming assays

Ten µl of 16 hour-old cultures of *P. mirabilis* strains (BL95, BL139 and BL141) were spotted on freshly poured LB agar plates (2 % agar). Plates were incubated at 37°C for 48 hours. Image is a representative of 3 replicates.

### Statistical analysis

Ordinary one-way ANOVA followed by multiple comparisons and Non-paired Welch T-Tests were performed where indicated using GraphPad Prism version 9.0.0 for MacOS, GraphPad Software, San Diego, California USA, www.graphpad.com.

### Data availability

All relevant data are within the manuscript and its supplemental material.

## ACKNOWLEDGEMENTS

Work was supported by UIC Institutional Startup Funds to JB and UIC Provost’s Graduate Research Award to DK.

The funders had no role in study design, data collection and interpretation, or the decision to submit the work for publication. The authors would like to thank Karine Gibbs for helpful discussions and for providing the *P. mirabilis* T6SS deficient strain.

The authors would also like to thank Henar Cuervo Grajal for her help and expertise in obtaining three-color fluorescence images. Plasmid pJC43 was a gift from Jean Celli, pMRE133 was a gift from Mitja Remus-Emsermann, pUCP30T-E2Crimson was a gift from Mariette Barbier, and p7INT-recA_luxAB was a gift from Michael Federle. *L. monocytogenes*, *K. pneumoniae* and *C. rodentium* were provided by Nancy Freitag and Donna MacDuff. Jason Devlin and Amisha Rana are acknowledged for their assistance with experiments, helpful discussions, and critical reading of the manuscript.

## ABBREVIATIONS

SPF: Specific pathogen free
T3SS: Type III secretion system
T4SS: Type IV secretion system
T5SS: Type V secretion system
T6SS: Type VI secretion system
Cdz: Contact-dependent inhibition by glycine zipper proteins
CDI: Contact-dependent inhibition
MccPDI: Microcin proximity-dependent inhibition
QS: Quorum sensing

---

**Supplemental Figure 1:**
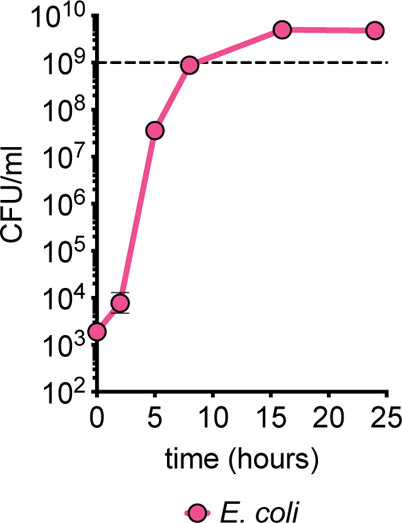
*E. coli* cells do not lose viability in a monoculture. *E. coli* cells were inoculated in LB and viability was measured over a 24-hour period. All data represent mean +/-SEM of at least three biological replicates. If error bars are not visible, error is smaller than symbol size.

**Supplemental Figure 2:**
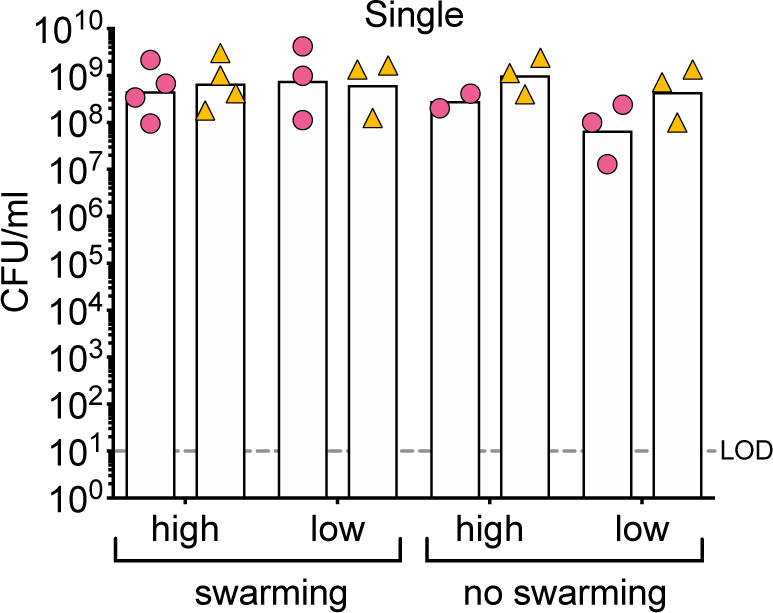
Cells remain viable in single cultures on solid surfaces. Monocultures of *P. mirabilis* and *E. coli* were inoculated on the solid surface of swarming permissive (LB) agar or non-swarming permissive (MacConkey) agar in high density (10^8^ CFU of each strain) or low density (10^3^ CFU of each strain) of cells in each spot. All data represent mean of at least three biological replicates (single high-density culture of *E. coli* on MacConkey two biological replicates). LOD = limit of detection

**Supplemental Figure 3:**
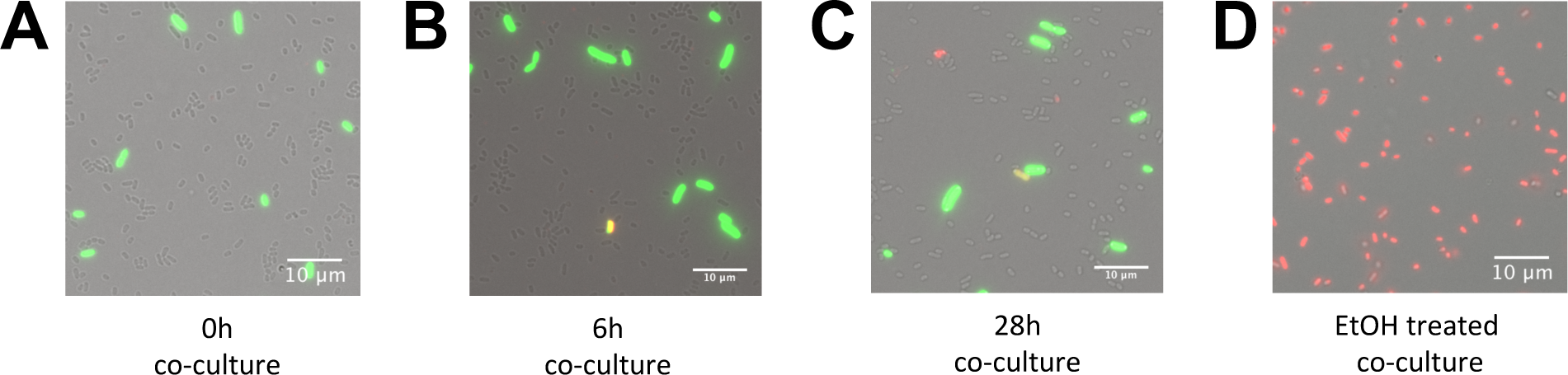
Killing does not cause loss of membrane integrity in target cells. Representative images of *P. mirabilis* (unstained, brightfield) and *E. coli* (GFP-positive) stained with propidium iodide (red) **(A)** at the start of co-culture (0 hour), **(B)** 6 hours and **(C)** 28h after the start of co-culture and **(D)** treated with 95% ethanol (EtOH).

**Supplemental Figure 4:**
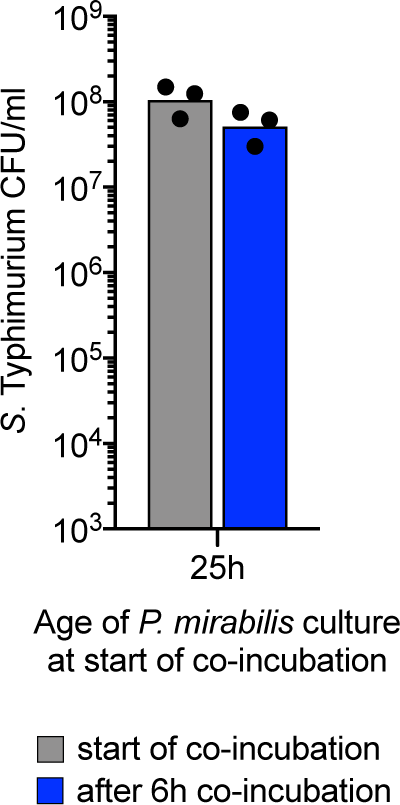
*P. mirabilis* in later stage of stationary phase does not kill *Salmonella enterica* serovar Typhimurium. *P. mirabilis* was grown in single culture for 25 hours before *S*. Typhimurium cells were added. Viability of *S*. Typhimurium was assessed at the beginning of co-culture and 6 hours after the start of co-culture.

